# An empirically-driven guide on using Bayes Factors for M/EEG decoding

**DOI:** 10.1101/2021.06.23.449663

**Authors:** Lina Teichmann, Denise Moerel, Chris Baker, Tijl Grootswagers

## Abstract

Bayes Factors can be used to provide quantifiable evidence for contrasting hypotheses and have thus become increasingly popular in cognitive science. However, Bayes Factors are rarely used to statistically assess the results of neuroimaging experiments. Here, we provide an empirically-driven guide on implementing Bayes Factors for time-series neural decoding results. Using real and simulated Magnetoencephalography (MEG) data, we examine how parameters such as the shape of the prior and data size affect Bayes Factors. Additionally, we discuss benefits Bayes Factors bring to analysing multivariate pattern analysis data and show how using Bayes Factors can be used instead or in addition to traditional frequentist approaches.

## 1. Introduction

The goal of multivariate decoding in cognitive neuroscience is to infer whether information is represented in the brain (Hebart & Baker, 2018). To draw meaningful conclusions in this information-based framework, we need to statistically assess whether the conditions of interest evoke different data patterns. In the context of time-resolved neuroimaging data, activation patterns are extracted across MEG or EEG sensors and classification accuracies are used to estimate information at every timepoint (see Figure 1 for an example). Currently, null hypothesis statistical testing (NHST) and p-values are the de-facto method of choice for statistically assessing classification accuracies, but recent studies have started using Bayes Factors (Grootswagers et al., 2021; e.g., Grootswagers, Robinson, & Carlson, 2019b; Grootswagers, Robinson, Shatek, et al., 2019; Kaiser et al., 2018; Karimi-Rouzbahani et al., 2021; Mai et al., 2019; Proklova et al., 2019; Robinson et al., 2019, 2021). Under the null hypothesis, the mean equals chance decoding and under the alternative hypothesis the mean is larger than chance decoding. The direct comparison of the predictions of two hypotheses is one of the strengths of the Bayesian framework of hypothesis testing (Jeffreys, 1939, 1935). Bayes Factors describe the probability of one hypothesis over the other given the observed data. In the multivariate pattern analysis (MVPA) context, this means we use Bayes Factors to test the probability of above-chance classification versus at-chance classification given the decoding results across participants at each timepoint. The goal of the current paper is to present and discuss Bayes Factors from a practical standpoint in the context of time-series decoding, while referring the reader to published work focusing on the theoretical and technical background of Bayes Factors.

**Figure 1.**
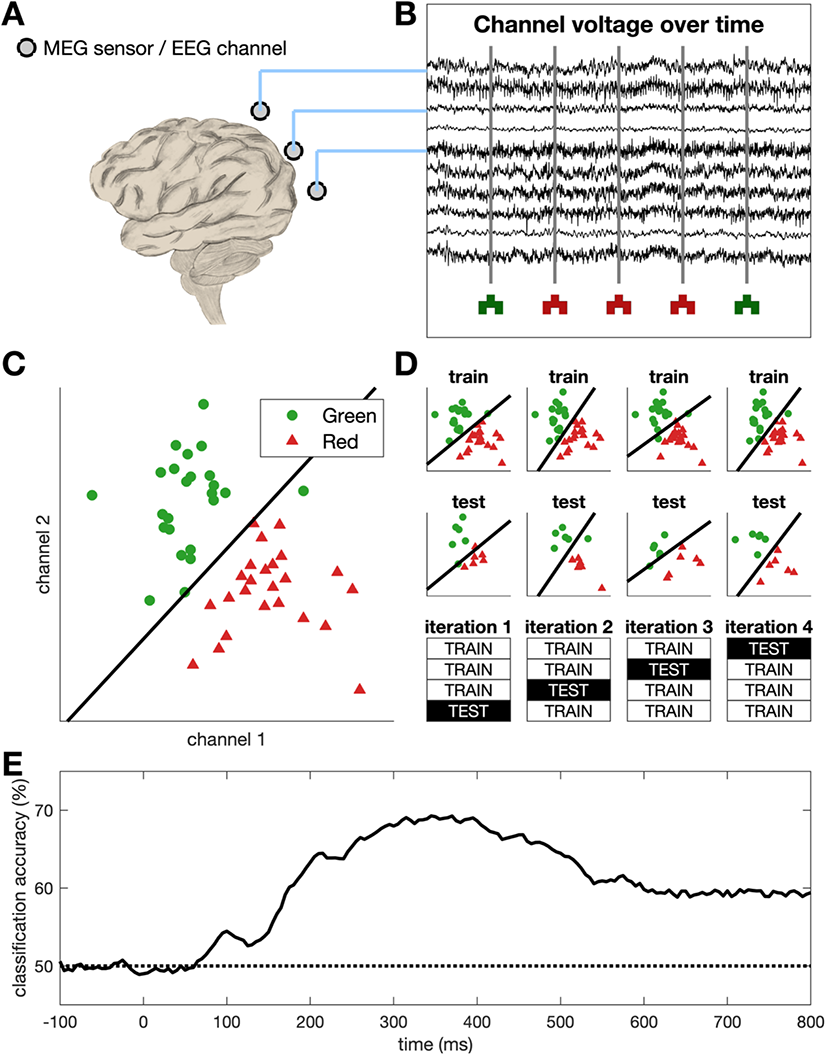
Overview of MVPA for time-series neural data. (A) Example MEG sensors / EEG channels. (B) Simulated time-series neuroimaging data for a few sensors/channels. Vertical lines show stimulus onsets with example stimuli plotted below. Data is first epoched from −100 to 800 ms relative to stimulus onset, resulting in multiple time-series chunks associated with seeing a red or a green shape. (C) Using the epoched data, we can extract the sensor/channel activation pattern across the different sensors/channels (only 2 displayed for simplicity) for every trial at every timepoint. Then a classifier (black line) is trained to differentiate between the activation patterns evoked by red and green trials. The shape of the stimuli is not relevant in this context. (D) Example of a 4-fold cross validation where the classifier is trained on three quarters of the data and tested on the left-out quarter. This process is repeated at every timepoint. (E) We can calculate how often the classifier accurately predicts the colour of the stimulus at each timepoint by averaging across all testing folds. Theoretical chance level is 50% as there are two conditions in the simulated data (red and green). During the period before stimulus onset, we expect decoding to be at chance, and thus the baseline period can serve as a sanity check.

The Bayesian approach brings several advantages over the traditional NHST framework (Dienes, 2011, 2014, 2016b; Keysers et al., 2020; Morey et al., 2016; Wagenmakers et al., 2018). In addition to allowing us to contrast evidence for above-chance versus at-chance decoding directly, Bayes Factors are a measure of strength of evidence for one hypothesis versus another. That means, we can *directly* assess how much evidence we have for different analyses. For example, if we were interested in testing whether viewing different colours evokes different neural responses, we could examine differences in the neural signal evoked by seeing red, green, and yellow objects. Using Bayes Factors, we could then directly compare whether red versus green can be decoded as well as red versus yellow. Larger Bayes Factors reflect more evidence which makes the interpretation of statistical results across analyses more intuitive. Another advantage is that Bayes Factors can be calculated iteratively while more data are being collected and that testing can be stopped when there is a sufficient amount of evidence (Keysers et al., 2020; Wagenmakers et al., 2018). Such stopping-rules could be accompanied by a pre-specified acquisition plan and potentially an (informal) preregistration via portals such as the Open Science Framework (Foster & Deardorff, 2017). Using the data to determine when enough evidence has been collected is particularly relevant for neuroimaging experiments, as it might significantly reduce research costs and reduce the risk of having underpowered studies. Thus, using a Bayesian approach to statistically assess time-series classification results can be beneficial both from a theoretical as well as an economic standpoint and might ease the ability to interpret and communicate scientific findings.

While Bayes Factors provide an alternative to the more traditional NHST framework, incorporating Bayes Factors into existing time-series decoding pipelines may seem daunting. Introductory papers often focus on mathematical aspects, and on relatively straightforward behavioural experiments (e.g., Ly et al., 2016; Morey et al., 2016; Rouder et al., 2009). We present an example based on a previously published time-series decoding study (Teichmann et al., 2019) and will present results from simulations to show the influence of certain parameters on Bayes Factors. We make use of the established Bayes Factor R package (Morey et al., 2015) to calculate the Bayes Factors but provide sample codes along with this paper showing how to access the Bayes Factor R package via Matlab and Python (https://github.com/LinaTeichmann1/BFF_repo). We also show how the Bayes Factors in our example compare to p-values. Based on empirical evidence, we will give recommendations for Bayesian analysis applied to M/EEG classification results. The aim of this paper is to provide a broad introduction to Bayes Factors from a viewpoint of time-series neuroimaging decoding. We aim to do so without going into the technical or mathematical detail, and instead provide pointers to relevant literature on the specifics.

## 2. Methods & Results

### 2.1 Example dataset & inferences based of Bayes Factors

The aim of the current paper is to show how to use Bayes Factors when assessing time-series neuroimaging classification results and test what effect different analysis parameters have on the results. We have used a practical example of previously published MEG data (Teichmann et al., 2019), which we re-analysed using Bayes Factors. In the original experiment, eighteen participants viewed coloured shapes and grayscale objects in separate blocks while the neural signal was recorded using MEG. Here, we only considered the coloured shape trials (“real colour blocks”, 1600 trials in total). Identical shapes were coloured in red or green and were shown for 100 ms followed by an inter-stimulus-interval of 800-1100 ms. The data was epoched from −100 ms to 800 ms (200 Hz resolution) relative to stimulus onset and a linear classifier was used to differentiate between the neural responses evoked by red and green shapes. A 5-fold cross-validation was used with the classifier being trained on 80% of the data and tested on the remaining 20%. This classification analysis resulted in decoding accuracies over time for each participant. In the original study, permutation tests and cluster-corrected p-values were used to assess decoding accuracies as implemented in CoSMoMVPA (Oosterhof et al., 2016). Here, we calculated Bayes Factors instead and examined how parameter changes affected the results.

When running statistical tests on classification results, we are interested in whether decoding accuracy is above-chance at each timepoint. To test this using a frequentist approach, we can use permutation tests to establish whether there is enough evidence to reject H_0_ which states that decoding is equal to chance. If there is enough evidence, we can reject H_0_ and conclude that decoding is different from chance. Given that below-chance decoding accuracies are not meaningful, we usually are interested only in above-chance decoding (directional hypothesis). In contrast to the frequentist approach, Bayes Factors quantify how much the plausibility of two hypotheses changes, given the data (see e.g., Ly et al., 2016). Here, we ran a Bayesian t-test of Bayes Factor R package (Morey et al., 2015) at each timepoint, testing whether the data is more consistent with H_a_ (decoding is larger than chance) over H_0_ (decoding is equal to chance). The resulting Bayes Factors center around 1 with numbers smaller than 1 representing evidence for H_0_ and numbers larger than 1 representing evidence for H_a_. In contrast to p-values, Bayes Factors are directly interpretable and comparable (cf. Keysers et al., 2020; Morey et al., 2016; Wagenmakers et al., 2016). That is, a Bayes Factor of 10 means the data is 10 times more likely to be observed under H_a_ as opposed to H_0_. Similarly, a Bayes Factor of 1/10 means the data is 10 times more likely to be observed under H_0_ as opposed to H_a_. Thus, in the context of time-series decoding, Bayes Factors allow us to directly assess whether and how much evidence there is at a given timepoint for the alternative over the null hypothesis and *vice versa* (Figure 2C).

**Figure 2.**
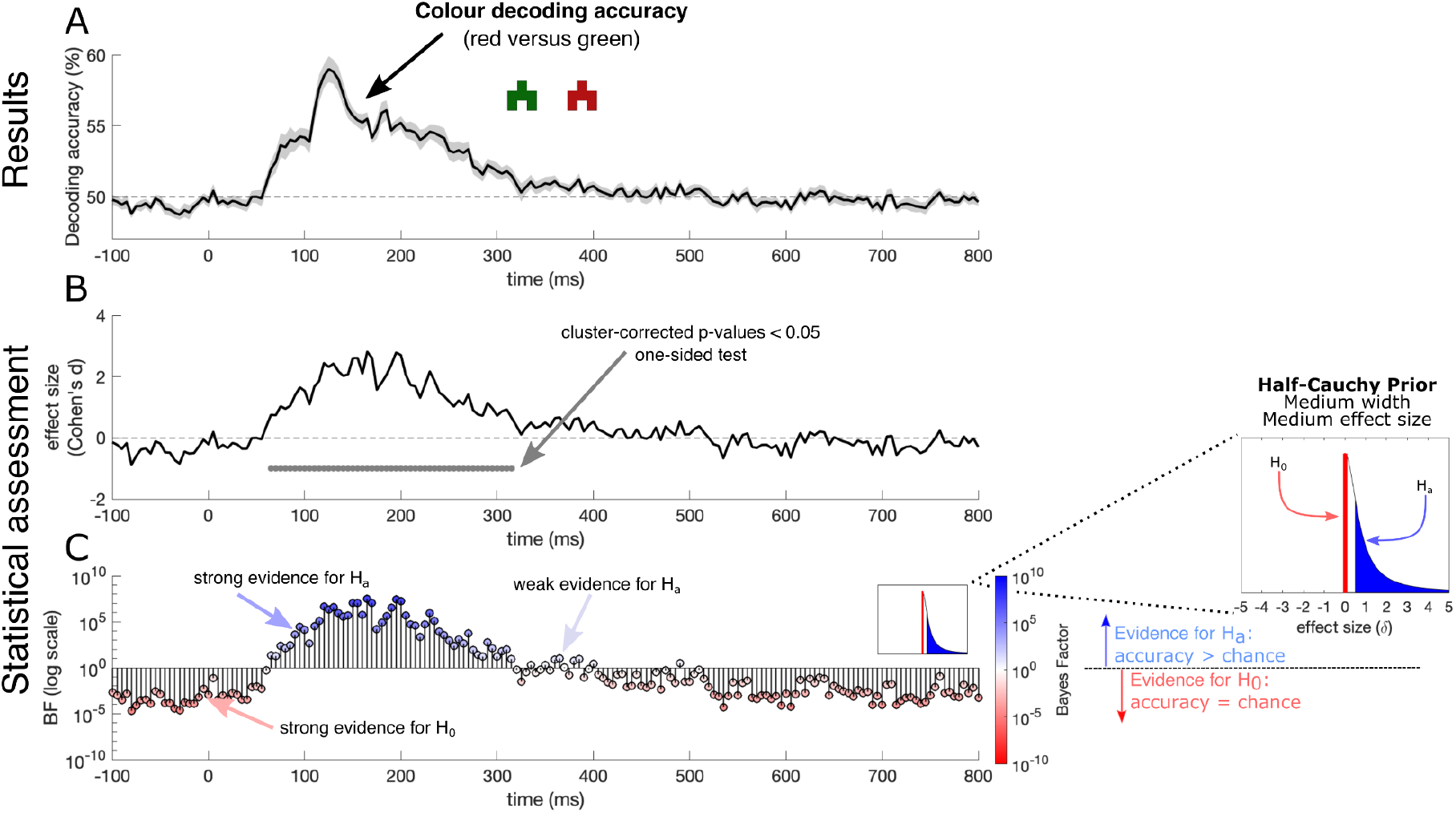
Decoding results of our practical example dataset with statistical assessments. (A) Colour decoding over time (black line). The dashed line shows theoretical chance decoding (50%). The grey shaded area represents the standard error across participants. (B) Effect size over time with the cluster-corrected p-values at each timepoint printed below in grey. (C) Bayes Factors over time for this dataset on a logarithmic scale. Blue, upwards pointing stems indicate evidence for above-chance decoding and red, downwards pointing stems show evidence for at-chance decoding at every timepoint. We used a hybrid one-sided model comparing evidence for above-chance decoding versus a point-nil at *δ* = 0 (no effect). For the alternative hypothesis, we used a half-Cauchy prior with medium width (r = 0.707) covering an interval from *δ* = 0.5 to *δ* = ∞. The half-Cauchy prior assumes that small effect sizes are more likely than large ones, but the addition of the interval deems very small effects *δ* < 0.5 as irrelevant. During the baseline period (i.e., before stimulus onset), the Bayes Factors strongly support the null hypothesis, confirming the sanity check expectation.

### 2.2 Adjusting the prior range to account observed chance decoding

Bayes Factors represent the plausibility that the data emerged from one hypothesis compared to another. In the example dataset, the two hypotheses are that decoding is at chance (i.e., H_0_, no colour information present) or that decoding is above chance (i.e., H_a_, colour information present). To deal with the fact that observed chance decoding can be different than the theoretical chance level, we can adjust the prior range of the alternative hypothesis to allow for small effects under the null hypothesis (Rouder et al., 2009). The prior range (called “null interval” in the R package) is defined in standardized effect sizes and consists of a lower and upper bound. To incorporate the differences between observed and theoretical chance level, we can define a range of relevant effect sizes for the alternative hypothesis, for example, from *δ* = 0.5 to *δ* = ∞. To determine which values are reasonable as the lower bound of this interval, we changed the prior range systematically and examined the effect on the resulting Bayes Factors (Figure 3). We found that smaller lower bounds at *δ* = 0 and *δ* = 0.2 resulted in weaker evidence supporting the null hypothesis than ranges starting at *δ* = 0.5 and *δ* = 0.8. The range did not have a large effect on timepoints with strong evidence for H_a_. The effect of changing the prior range is larger for the null hypothesis than the alternative as chance decoding is not exactly 50% but distributed around chance. Changing the lower bound of the prior range means that the effects that are just larger than *δ* = 0 can support the null hypothesis. Thus, the results here demonstrate that we can compensate for the differences between theoretical and observed chance by adjusting the prior range and effectively considering small effect sizes as evidence for the null hypothesis rather than the alternative.

**Figure 3.**
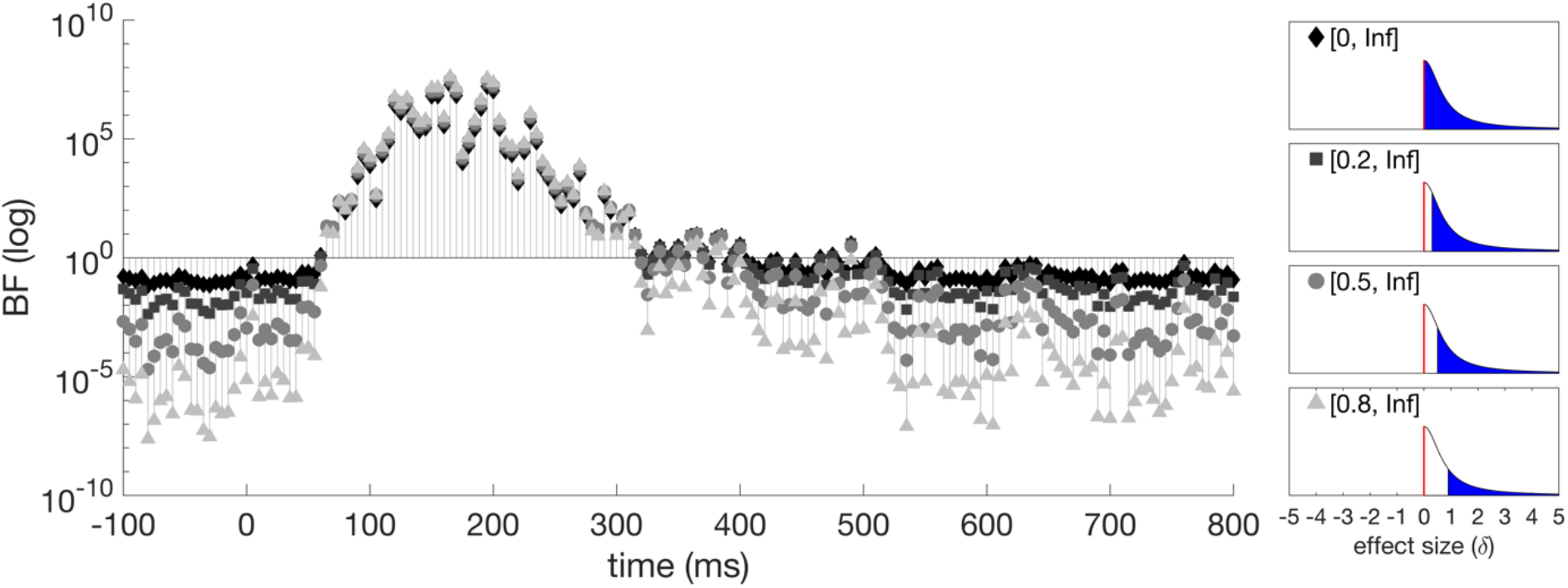
The effect of changing the prior range (null interval) on Bayes Factors in our example data. Intervals starting at larger effect sizes led to more timepoints showing conclusive evidence for H_0_. This is due to the fact that theoretical and observed chance levels are not the same. The panels on the right show the prior distributions with the different null intervals.

To further examine what a reasonable lower bound of the prior range is, we looked at effect sizes observed during the baseline window (before stimulus onset) in a selection of our previous studies (Grootswagers et al., 2021; Grootswagers, Robinson, & Carlson, 2019a; Moerel, Grootswagers, et al., 2021; Moerel, Rich, et al., 2021; Teichmann et al., 2018, 2020). Using the baseline window allows us to quantify the difference between theoretical and observed chance, as we do not expect any meaningful effects before stimulus onset (e.g., stimulus colour is not decodable before the stimulus is presented). Thus, the baseline period can effectively tell us which effect sizes can be expected by chance. Using this method, we estimated maximum effect sizes for different analyses in each paper (see different bars in Figure 4). Across our selection of prior studies, we found an average maximum effect size of *δ* = 0.39 before stimulus onset and an average maximum effect size of *δ* = 1.91 after stimulus onset (Figure 4). This survey shows that effect sizes as large as *δ* = 0.5 can be observed when no meaningful information is in the signal. Thus, this supports the conclusions from the example dataset showing that prior ranges with a lower bound of *δ* = 0.5 may be a sensible choice when using Bayes Factors to examine time-series M/EEG decoding results.

**Figure 4.**
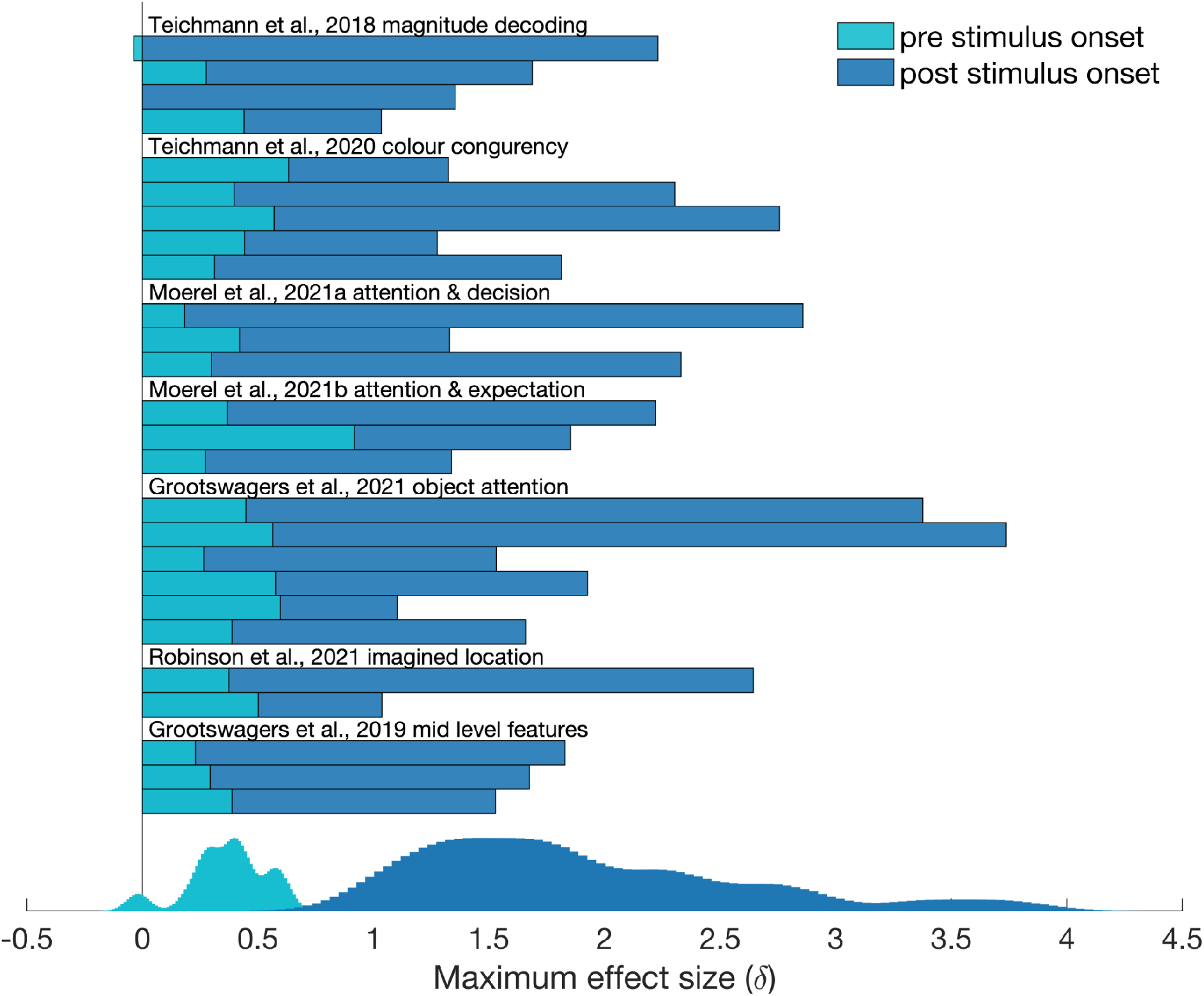
Estimated maximum effect sizes during baseline and after stimulus onset for prior decoding studies that used visual stimuli. Using already published data, we calculated the maximum effect sizes during the baseline (light blue) and post-stimulus (dark blue) to estimate typical peak effect sizes in visual decoding studies. Each bar represents a unique analysis within the paper. The estimations show that a reasonable range for H_a_ would start at *δ* = 0.5 or above, as during baseline decoding accuracies corresponding to standardized effect sizes as high as *δ* = 0.5 were observed.

### 2.3 Changing the prior width to capture different effect sizes

Another feature that can be changed in the Bayesian t-test is the width of the half-Cauchy distribution (referred to as r-value in the Bayes Factor Package). Small r-values create a narrower, sharply peaking distribution, whereas larger values make the distribution wider with a prolonged peak. Standard prior widths incorporated in the Bayes Factor R package are medium (r = 0.707), wide (r = 1), and ultrawide (r = 1.414). Keeping the prior range consistent ([0.5, Inf]) while using the three prior widths implemented into the R Bayes Factor Package (medium = 0.707; wide = 1; ultrawide = 1.414). We found that changing the width of the Cauchy prior did not have a pronounced effect on the Bayes Factors (Figure 5). In our specific example, this is probably the case because the effect sizes quickly rose to *δ* > 2 (Figure 2b) which means that the subtle differences between the different prior widths do not have a substantial effect on the likelihood of the data arising from H_a_ over H_0_. Thus, using the default prior width (r = 0.707) for the decoding context seems like a reasonable choice.

**Figure 5.**
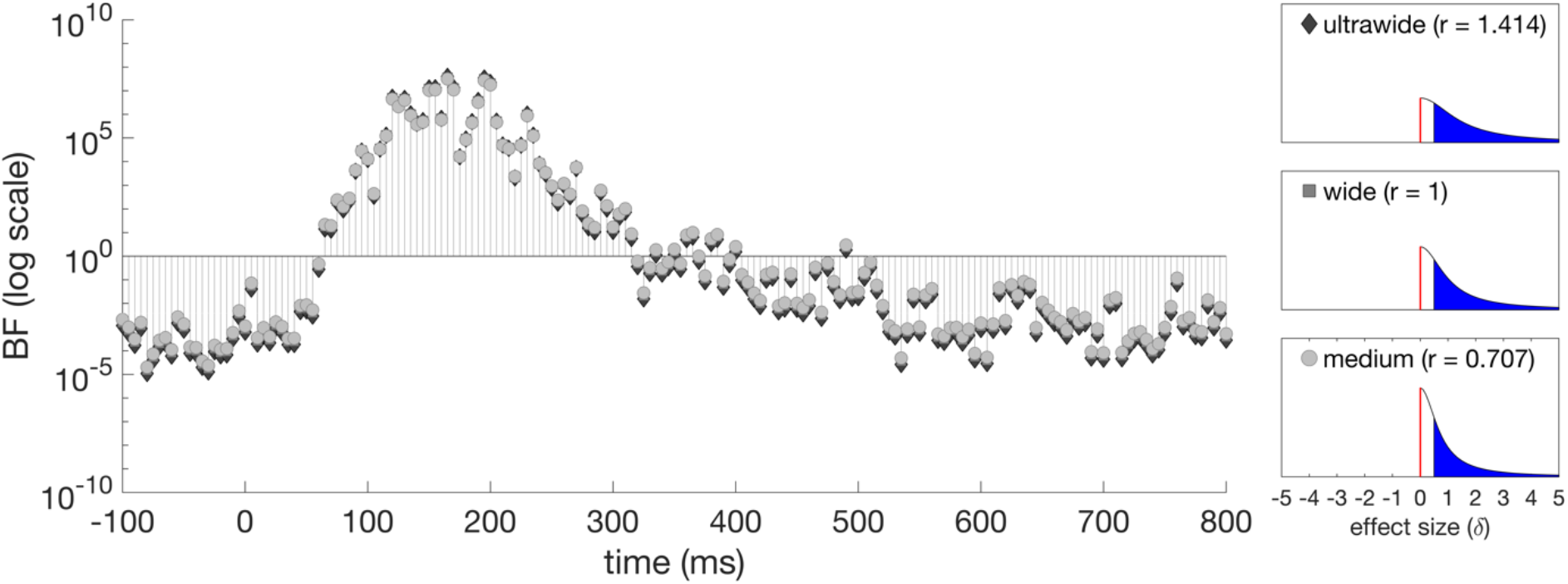
Bayes Factors over time for the example data set when the prior width is changed. The width of the prior had no pronounced effect on the Bayes Factors we calculated. The panels on the right show the prior distributions with the different widths.

### 2.4 The effect of data size on statistical inferences

In a lot of cases, there are financial and time limits on how many participants can be tested and for how long. To obtain an estimate of how much data is needed to draw conclusions and avoid ending up with underpowered studies, we used the example dataset and reduced the data size for analysis. As classification analyses are usually run at the subject level but statistical assessment is run at the group level, we tested how changing data size both by trial numbers and participant numbers influences Bayes Factors in the time-series decoding context (Figure 6). In the original example dataset, the classifier was trained on 1408 trials and tested on 352 trials (5-fold cross-validation). There were five different shapes in the red and the green condition (160 repetitions for each coloured shape) and the cross-validation schema was based on leaving all trials of one shape out for testing. Statistical inferences were drawn on the group level which contained data from 18 participants. To examine the effect of data size (and effectively noise level) on the Bayes Factor calculations, we re-ran the analysis reducing the data size first by retaining the first 1200 (75%), 800 (50%), 400 (25%), or 160 (10%) trials participants completed. We cross-validated in the same way as in the original paper, with the only difference being how many trials of each shape were included. In addition, we subsampled from the whole group, retaining data from the first 6, 12, or all 18 participants and re-ran the statistical analysis. We then compared the results from the reduced-size colour datasets using Bayes Factors and cluster-corrected p-values^1^.

**Figure 6.**
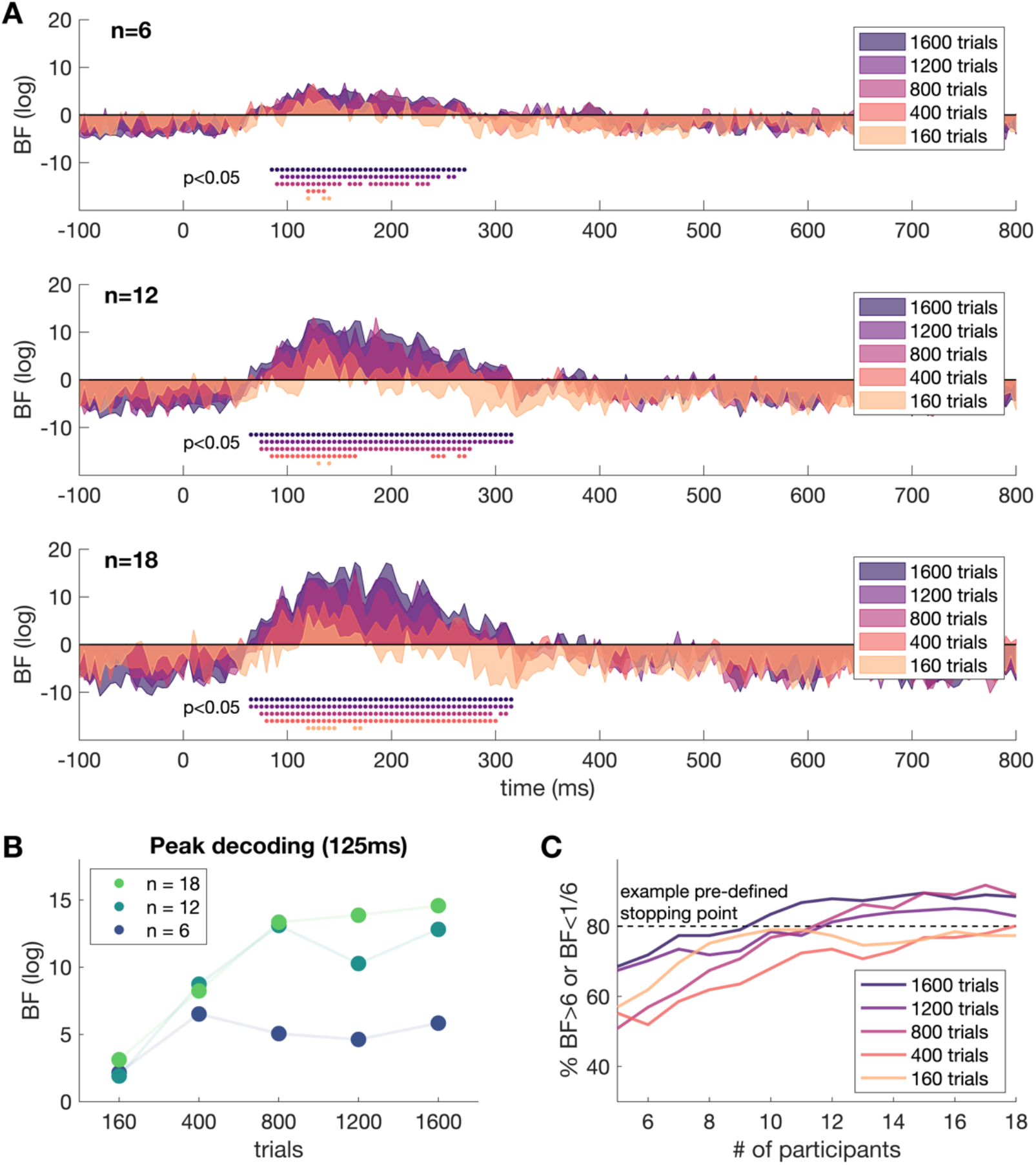
Results of the colour MEG decoding study, using a limited number of trials and participant data to simulate a piloting scenario. (A) The first three plots show Bayes Factors over time along with cluster-corrected p-values. The colour in all plots reflects the number of trials used to train and test the classifier. (B) Compares Bayes Factors at peak decoding (125ms) for the different data sizes. (C) Compares how many participants would have needed to be tested given the different number of trials with an example pre-defined stopping point. For example, with 1600 trials and >9 participants, 80% of the Bayes factors (at different time points) exceeded 6 or 1/6. With fewer trials, more participants are needed to reach this example stopping point.

Overall, our analyses highlight that we need to have a large enough number of trials and a large enough number of participants to draw firm conclusions about our time-resolved decoding results. Testing more participants resulted in stronger evidence for H_a_ and H_0_, with fewer timepoints in the inconclusive range (Bayes Factors) and more significant above-chance decoding timepoints (p-values). Similarly, running the classification with more trials, led to more timepoints with large Bayes Factors supporting H_a_ and more above-chance decoding timepoints. However, one of the key advantages of using Bayes Factors instead of p-values is that we can potentially obtain a good idea of how many trials are needed even if we run a pilot experiment with a limited number of participants. A reasonable strategy would be to overpower the subject-level data (i.e., number of trials) for the pilot sample and then sub-sample to explore how many trials are needed. In our example, we can see that the amount of evidence for H_a_ at peak decoding is not sufficient when we only use 160 trials (10% of the original sample), regardless of the number of subjects. Increasing the trials to 400 or 800 (25% or 50% of the original sample) leads to similar conclusions as using all 1600 trials. As Bayesian statistics allow for sequential sampling, we could collect data from more participants until a criterion is reached. For example, if we had pre-defined a stopping criterion as 80% of the timepoints being in the conclusive range (Bayes Factors larger than 6 or smaller than 1/6), we would have been able to stop collecting data after 9 participants completed 1600 trials or after 18 participants completed 400 (Figure 6c). Overall, the data suggest that insufficient data at the subject-level ultimately leads to inconclusive evidence, highlighting that a large number of trials is just as, if not more important, than large numbers of participants.

The example dataset provides insight into the effect of parameters such as data size and prior shape on Bayes Factors. However, it is possible that different studies find different effect sizes. We simulated larger datasets with fixed effect sizes between *δ* = 0 and *δ* = 1 to examine the interaction of sample size with different prior ranges for different effect sizes (Figure 7). We simulated 1000 datasets with specific effect sizes for each sample size and calculated the Bayes Factors. We then calculated the median Bayes Factor for each sample- and effect size combination to show how prior range choices interact with the possibility of finding evidence for effects of different sizes. Specifically, we compared a prior range of 0.5 to infinity (Figure 7A) to a prior range of zero to infinity (Figure 7B).

**Figure 7.**
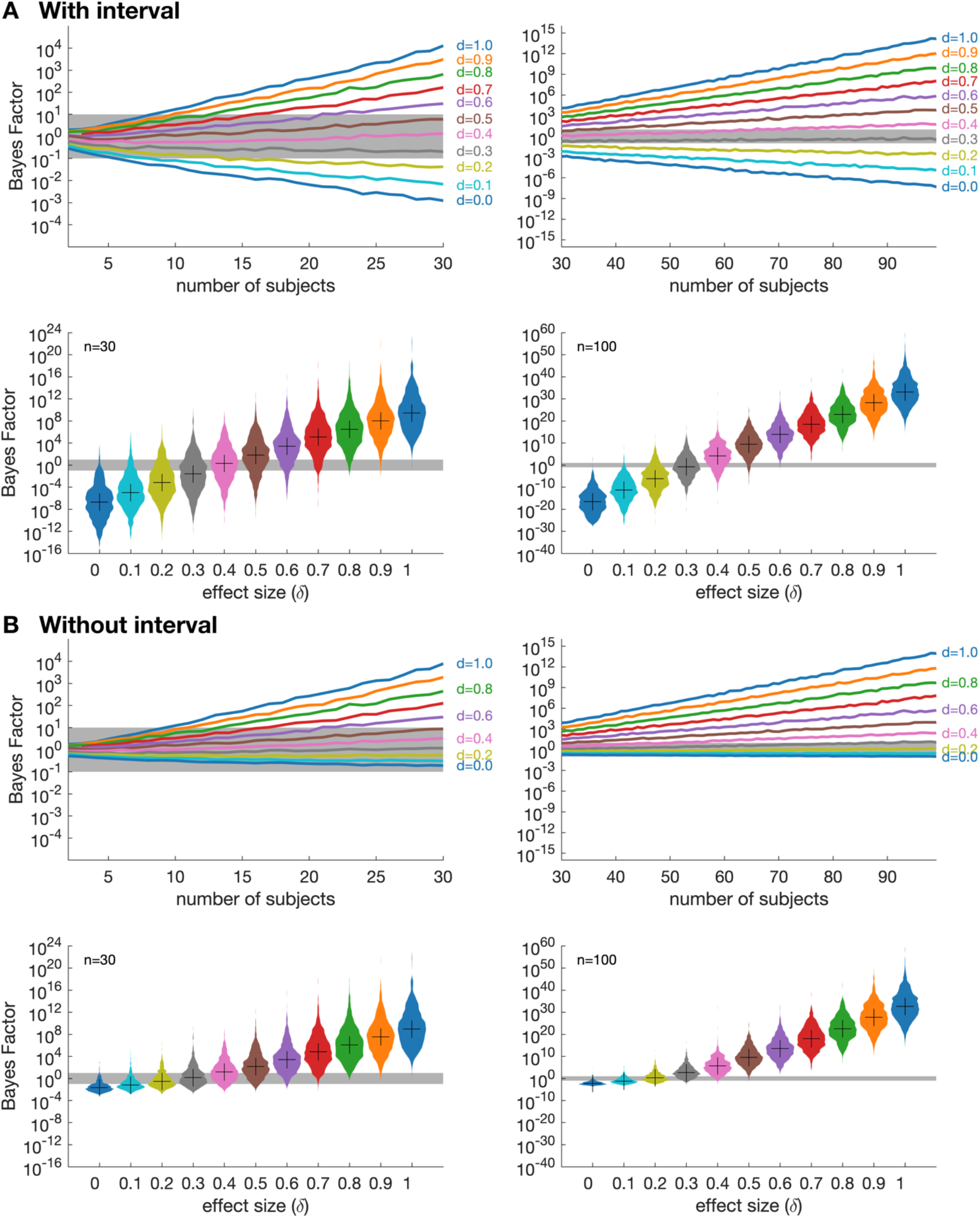
Simulated data varying effect sizes and numbers of participants highlight the rationale for using an interval. We performed 1000 simulations to demonstrate how the Bayes Factors behave with different sample sizes given different effect sizes. A shows Bayes Factors obtained by using a half-Cauchy prior with an interval [0.5 Inf]. B shows Bayes Factors obtained by using a half-Cauchy prior without an interval. The first and third rows show the median Bayes Factors of 1000 simulations as a function of the number of participants. The second and fourth rows show the distribution of the Bayes Factors from 1000 simulations using 30 participants (left panels) and 100 participants (right panels). The distributions of the Bayes Factors highlight the rationale for using an interval, as without an interval it is nearly impossible to find substantial evidence for the null hypothesis even when the effect size equals zero.

When specifying the prior range to 0.5 to infinity (Figure 7A), our results show that small sample sizes are sufficient to draw solid conclusions when the effect sizes are near the extremes. For example, the simulations showed that there is substantial evidence for H_0_ from a small sample size if the true effect is very small. In contrast, if the effect size fell in between the specified ranges for the prior of H_a_ and H_0_ (i.e., between 0 and 0.5), we found that small sample sizes tended to result in inconclusive Bayes Factors neither supporting H_a_ or H_0_. However, if the sample size increased, the confidence that these effects were “real” also increased and therefore resulted in stronger confidence supporting one of the hypotheses. Importantly, however, large sample sizes did not automatically lead to an interpretable Bayes Factor if the effect was truly in between the specified prior ranges of H_a_ and H_0_, indicating that sample size had no effect on Bayes Factors in this case.

Consistent with our results for the example data, the simulations also showed that changing the range of the prior has a strong effect on finding substantial evidence for H_0_. If the prior range for the alternative is specified to start at zero (Figure 7B), it was almost impossible to find any evidence for H_0_, even if the effect size was truly zero. Thus, the simulations show that defining the prior range with a gap between effects expected under H_0_ and H_a_ is critical and that more data leads to larger Bayes Factors, but only if there is a true underlying effect.

## Discussion

Bayes Factors have seen a recent increase in popularity in cognitive science, as they can be used to provide quantifiable evidence for contrasting hypotheses. However, their uptake has to date been slow for neuroimaging experiments. To facilitate their adoption, we have provided an empirically-driven guide on implementing Bayes Factors for time-series neuroimaging decoding, using both real and simulated data. We showed that using Bayes Factors and cluster-corrected p-values lead to similar results when statistically assessing time-series neuroimaging decoding results. However, the key advantages of using Bayes Factors are the ability to compare evidence for H_a_ with evidence for H_0_ and having results that are quantifiable (e.g., Dienes, 2014; Wagenmakers et al., 2016). Our results show that for time-series decoding data, half-Cauchy priors with default width and an interval ranging from effect sizes of 0.5 to infinity provide sensible results. We also show that even a small number of participants can yield informative Bayes Factors, which can be useful for making decisions on experimental design parameters (e.g., number of trials) during piloting stages of a study.

Our results showed that the overall conclusions derived from Bayes Factors and p-values were quite similar, highlighting that theoretical considerations should be the deciding factor when choosing a statistical approach to analyse neural time-series data. In the decoding context, p-values afford a dichotomous decision of whether there is enough evidence to reject the hypothesis that decoding is at chance at a given timepoint. Rejecting the null hypothesis is decoupled from any prior beliefs or theories (Dienes, 2011) and is linked to an accepted overall error rate such as α = 0.05. P-values allow us to test for the presence of an effect at a given timepoint using widely accepted thresholds for evidence. While Bayes Factors can in principle also be thresholded to draw dichotomous conclusions, one of the added benefits of Bayes Factors over p-values is the ability to quantify the evidence. Another useful benefit of using Bayes Factors to analyse time-series decoding data is that Bayes Factors allow us to accrue evidence for above-chance as well as at-chance decoding. For time-series analyses in particular, this is a useful feature as the time period prior to stimulus onset can be considered as a control period where we would expect evidence for the null hypothesis. Testing both hypotheses simultaneously can also be a beneficial feature when the research question involves hypotheses predicting certain time-periods without any information in the neural signal (e.g., “X happens before Y” versus “Y happens before X”). Thus, depending on the research question it may be clear which statistical approach suits the time-series decoding analysis best. Otherwise, as overall conclusions do not differ, Bayes Factors and p-values can be used in a complementary way to provide quantifiable evidence for and against the tested hypotheses as well as definitive decisions (see also Lakens et al., 2020; van Dongen et al., 2019; Wagenmakers et al., 2018).

Through our results, we provide an empirical, straightforward guide to help implement Bayes Factors and demonstrate the extent of practical benefits when using Bayes Factors for time-series neural decoding. Using a data-driven approach, we showed which analysis parameters are most suitable for statistical assessment of time-series decoding data with Bayes Factors. While the Bayes Factors in our example MEG decoding dataset were robust against changes in the predefined width of the prior, defining the prior range so that there is a gap between H_a_ and H_0_ was critical for finding evidence for the H_0_. This strong effect of the prior range on the resulting Bayes Factors is particularly relevant in the decoding context, as classification accuracies under the null are not symmetrically distributed around chance (cf. Allefeld et al., 2016). Thus, a gap between H_0_ and the lower bound of H_a_ ensures that small above-chance classification accuracies are not treated as evidence for H_a_. Furthermore, we systematically varied dataset size and showed that using Bayes Factors for time-series decoding data is particularly beneficial when there is limited, noisy data such as in a piloting scenario, as quantifiable evidence for one hypothesis over another gives a stronger sense of whether it is worth pursuing the research question with the piloted design, or make changes (e.g., modify trial numbers or add/remove conditions). Finally, Bayes Factors can be calculated sequentially while evidence accumulation is monitored to stop once a criterion is reached (Dienes, 2011; Rouder, 2014), which can save resources and avoid underpowered studies (Wagenmakers et al., 2018). One possibility is to define a stopping criterion in terms of a percentage of timepoints where evidence is in the conclusive range of Bayes Factors (e.g., 80% of Bayes Factors are above 6 or below 1/6). As longer baselines can artificially increase the percentage of conclusive timepoints, only timepoints after stimulus onset should be considered or the duration of the baseline period should be pre-defined. As researchers generally do not have unlimited resources, it is possible to also pre-register an upper limit for the sample size (e.g., maximum 50 participants).

An open question is to what extent our parameter choices generalize to different paradigms, analysis approaches, and modalities. The Bayes Factor parameters used here were optimized for time-series decoding. It is in principle possible to use Bayes Factors in a similar way to analyse other time-series data such as event related potentials, oscillations or regressions, however, the Bayes Factor parameters might have to be adjusted. Similarly, the analysis pipeline discussed here could be extended to other neural decoding modalities such as fMRI (see e.g., Moerel, Rich, et al., 2021). Pilot data or analyses of previous data can be used to examine how parameters have to be modified in order to get sensible results.

A final consideration is the multiple comparisons problem arising from statistically testing many time points. When using Bayes Factors, as long as the evidence for each hypothesis is interpreted at face value (and not thresholded for ‘significance’), we do not need to control for multiple comparisons (Dienes, 2011, 2016a; Świątkowski & Carrier, 2020). That is because once we have established a prior and collected the data, we examine how much we have to adjust our prior beliefs given the data and compare the adjustment required for both hypotheses. This idea is not related to overall error rates and thus does not change if we sample data sequentially or run multiple tests (Dienes, 2016a). If a research question strongly depends on a dichotomous decision on multiple tests, then we advise to report corrected p-values (for which correction methods are well established) alongside the Bayes Factors.

In conclusion, we have provided an empirically-driven guide on how to use and interpret Bayes Factors for time-series neuroimaging decoding data. We show that Bayes Factors bring several advantages to interpreting time-series decoding results such as quantifiable evidence and an ability to compare evidence for above-chance with evidence for at-chance decoding. We hope this guide, and the accompanying example code (https://github.com/LinaTeichmann1/BFF_repo) can serve as a starting point to incorporate Bayesian statistics to existing analysis pipelines.

## Acknowledgements

This research was supported (in part) by the Intramural Research Program of the NIMH (ZIAMH002909). We thank Lincoln Colling for contributing to the repository.

## Footnotes

1 In comparison to the original paper, we did not use trial label permutations. Instead, we performed sign-flip permutations (which reduces the computational time) as implemented in CoSMoMVPA to generate the null distribution.

## Notes

### Competing Interest Statement

The authors have declared no competing interest.

### Summary of Updates

Figure 5 and Figure 3 were accidentally swapped in the last version

